## Supplementary material for "Inferring a simple mechanism for alpha-blocking by fitting a neural population model to EEG spectra": S1 Appendix

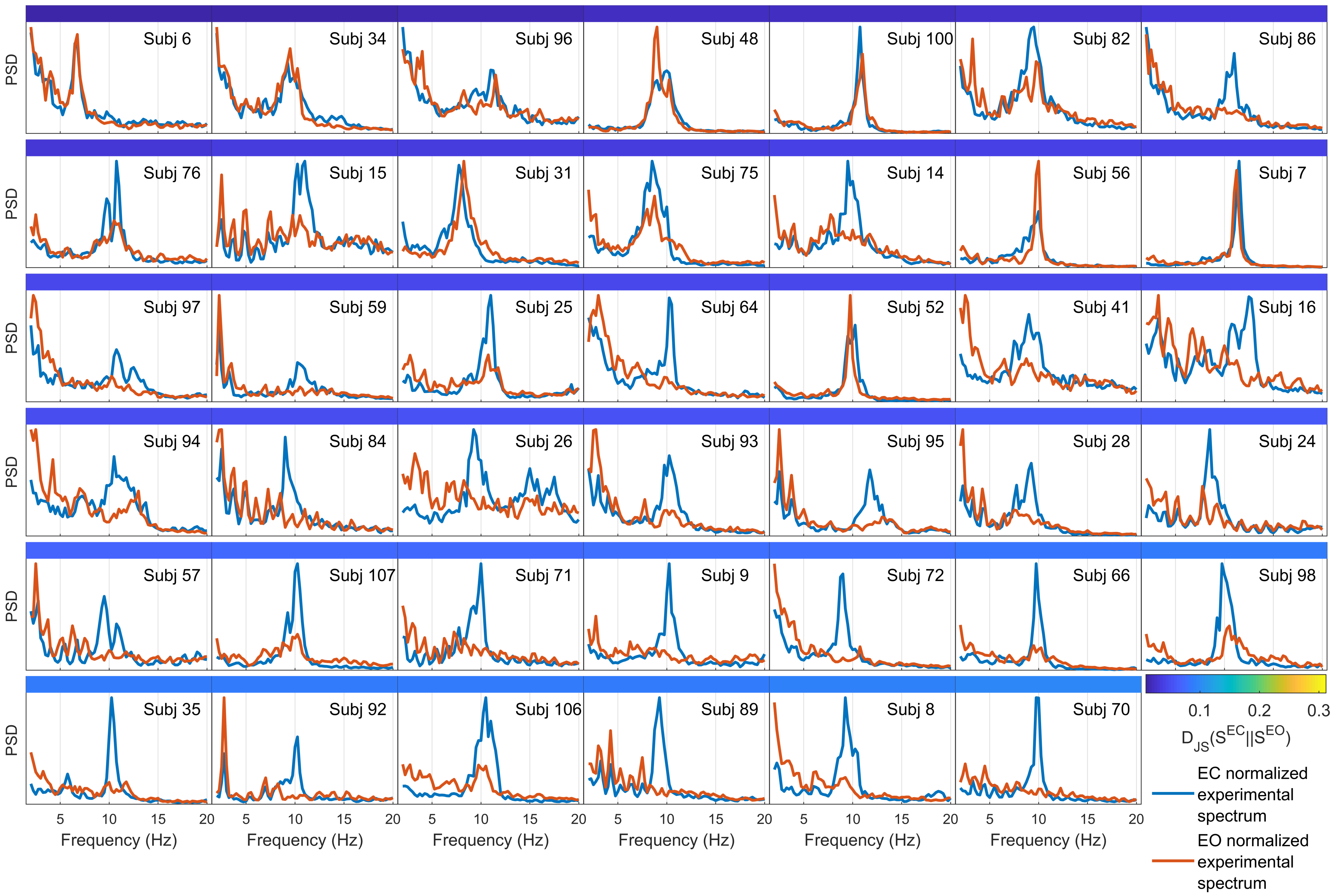


**Fig A$^{(i)}$. Degree of alpha-blocking across all subjects$^{(i)}$.** Shown are the pairs of EC and EO spectra across the 82 subjects. Subjects are ordered by degree of alpha blocking, as measured by the Jensen-Shannon divergence between the two states.

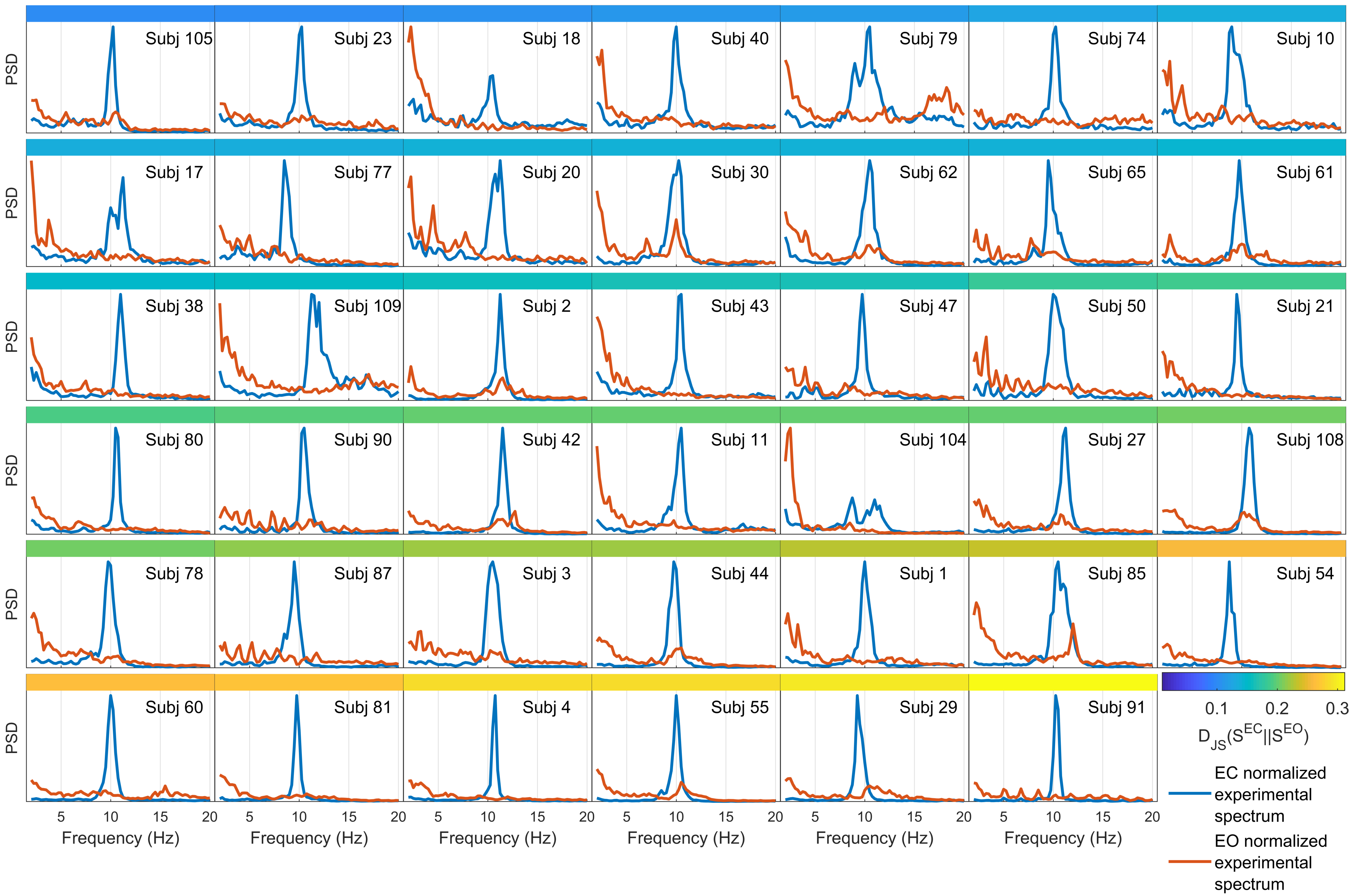


**Fig A$^{(ii)}$. Degree of alpha-blocking across all subjects$^{(ii)}$.**

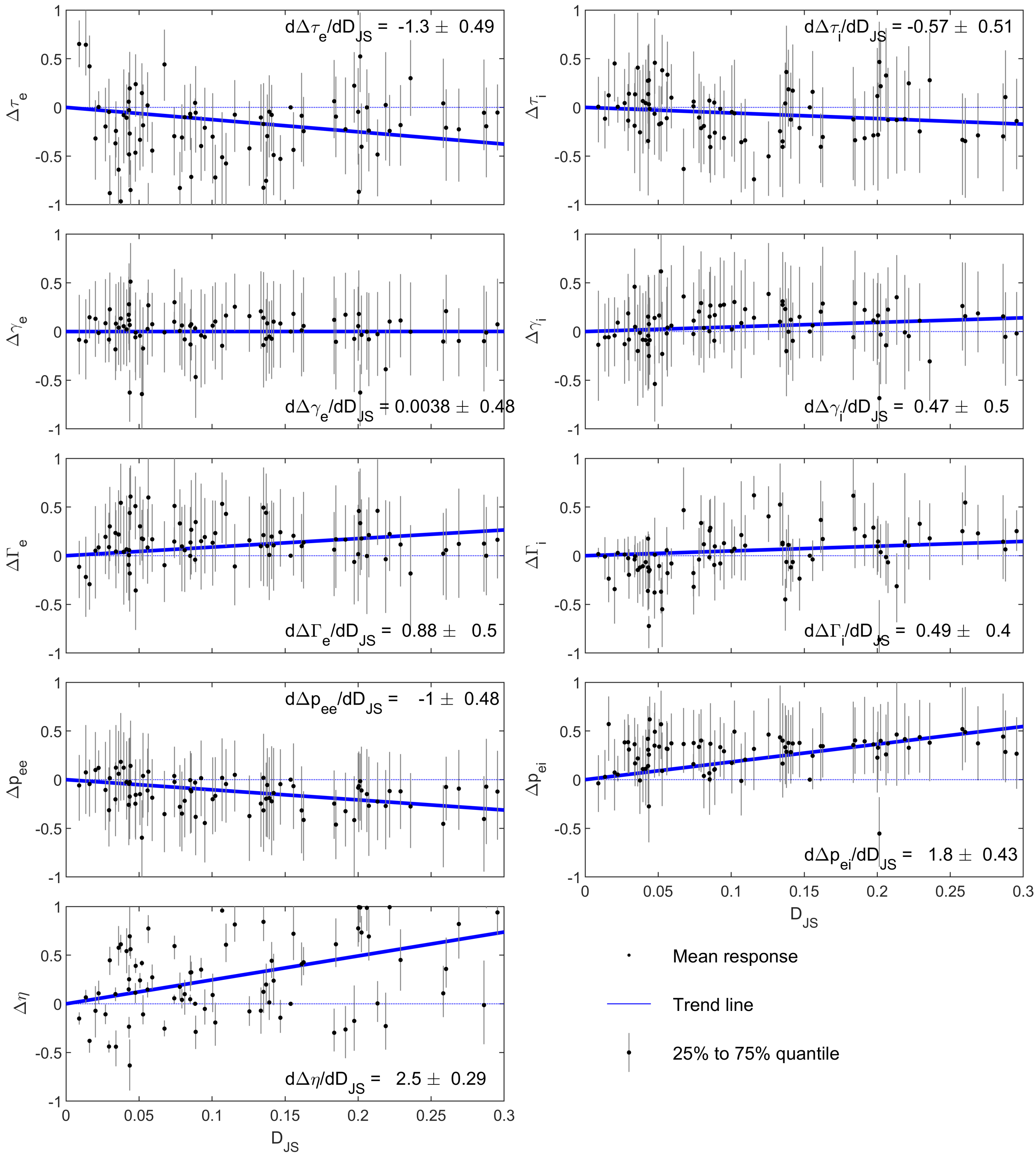


**Fig B. Unregularized EC to EO parameter responses and how they scale with the degree of alpha blocking.** The plots in this figure are similar to those in Fig 4 in the main article, except here the fitting was not regularized ($\lambda = 0$ in Eq 5, main paper). Considerably higher uncertainties are found in the parameter response estimates, making it difficult to observe whether some parameters are more important than others in driving alpha blocking. Note that the y-axis span is now 2 units, compared with 0.7 units for Fig 4, to accommodate the larger variability.

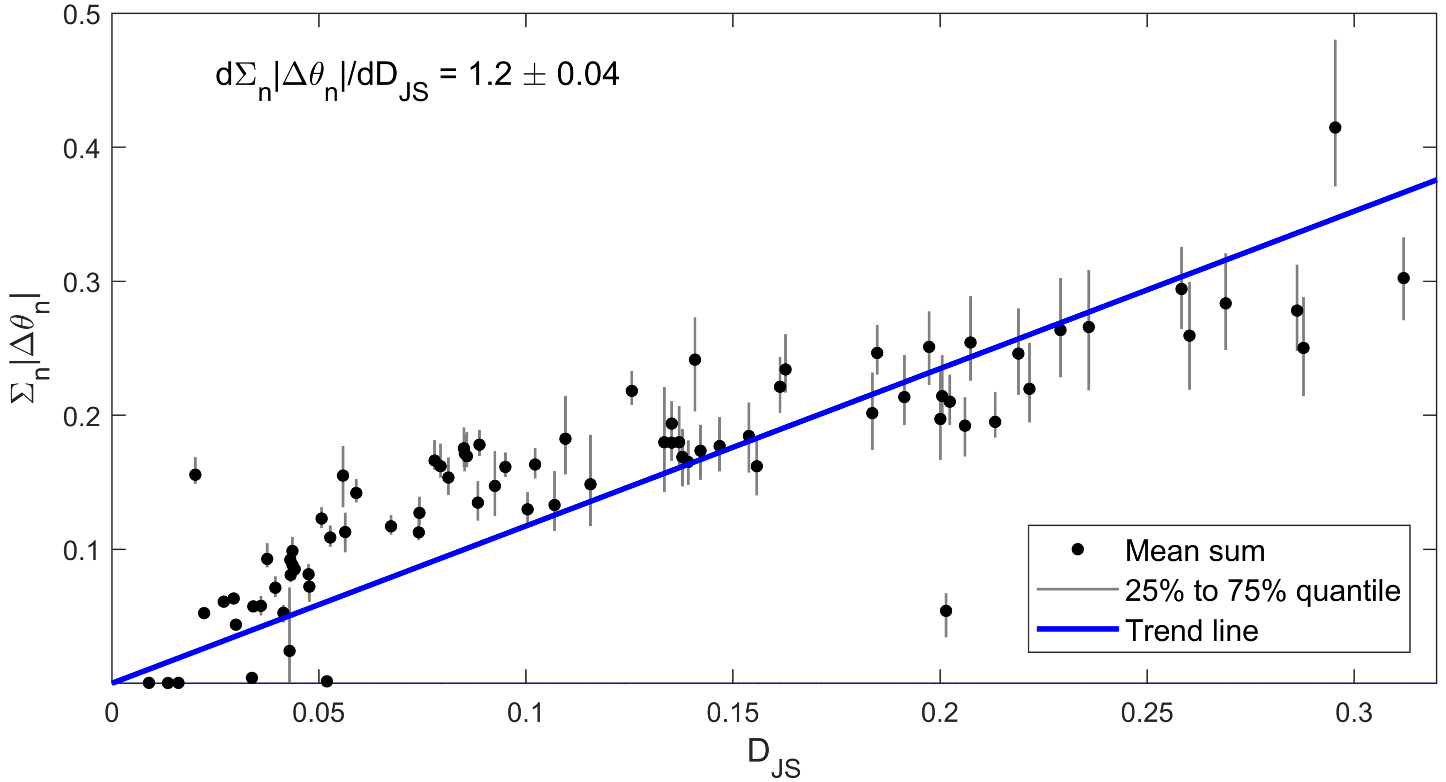


**Fig C. Manhattan distances between EC and EO parameter sets as a function of the degree of alpha blocking.** For each sampled parameter set, the Manhattan distance between the EC normalized parameter set and the EO normalized parameter set is computed from $\sum_m |\theta_m|$, where $m$ is a state-distinct parameter index. A black dot represents the mean of the 100 fitting samples found for each subject while error bars represent the interquartile range of Manhattan distances for each subject. A linear trend line with zero intercept is shown. The gradient $d\Sigma_m|\Delta\theta_m|/dD_{JS}$ and its uncertainty are computed based on $10^5$ random samples drawn from the 82 Manhattan distance distributions. The data show that the Manhattan distance in parameter space closely tracks the Jensen-Shannon divergence in spectra space, indicating that model response scales with the degree of alpha blocking and that the combined parameter response is identifiable.

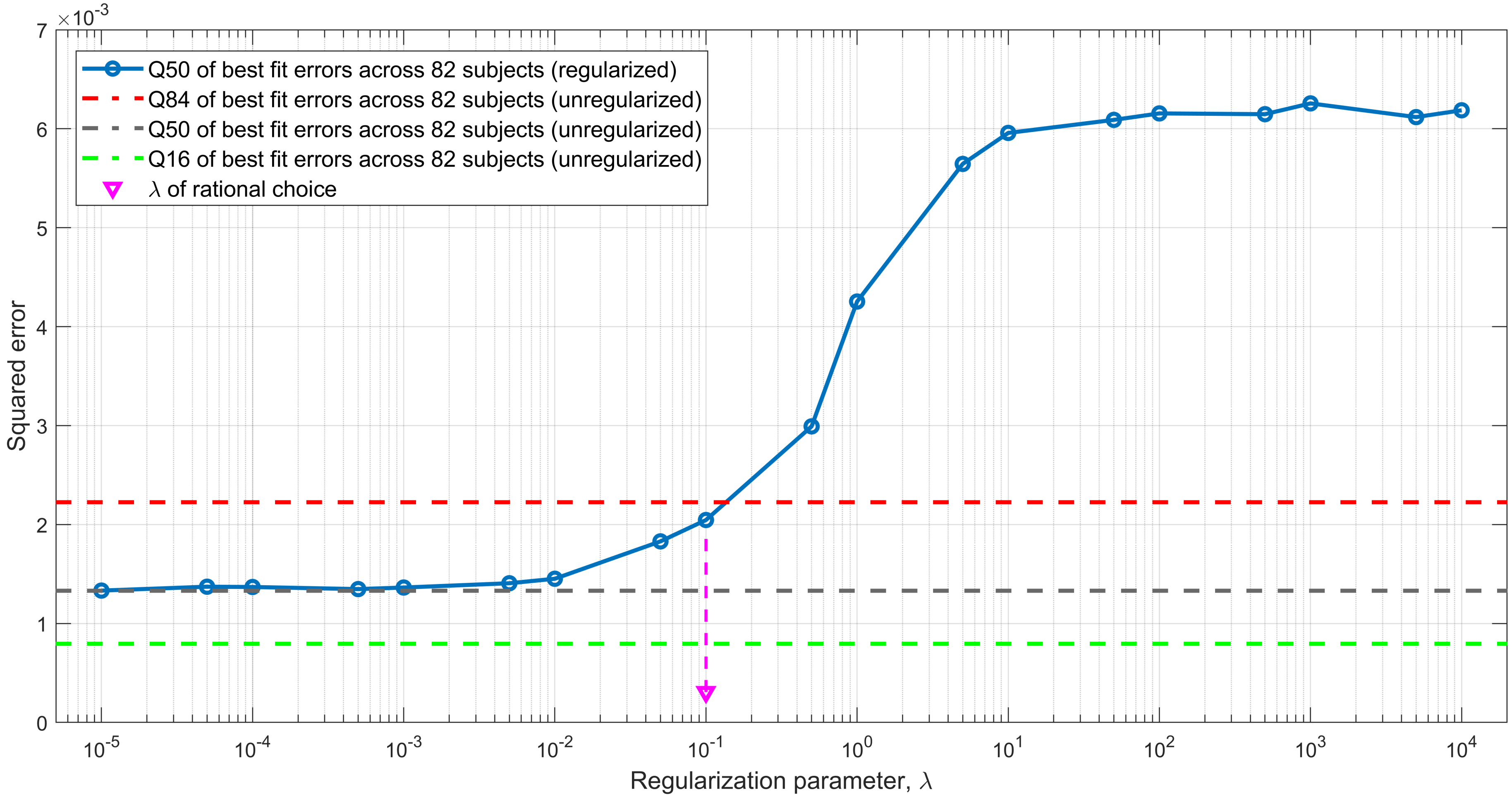


**Fig D. Comparison of fitting error as a function of regularization parameter** The regularization parameter, $\lambda$, represents the degree to which the parameter responses are penalized, with higher $\lambda$ meaning more penalization. As expected, the median least-squared errors (calculated over all 82 subjects) increases with $\lambda$, eventually reaching saturation when the parameter responses are forced to zero and the same prediction is made for both EC and EO. Shown as a benchmark are the 16%, 50%, and 84% quantiles of the fitting errors from the unregularized case. We choose the optimal $\lambda$ to be when errors reach the 84% quantile ($\lambda = 0.1$). At this optimum, many of the parameter responses have been forced to zero (see Fig 4, main paper) even though the spectral fits are still visually close to those for the unregularized case (see Fig 2, main paper).
